# CRISPR/Cas9 targeted genetic screening in Physcomitrella identifies novel cell division genes

**DOI:** 10.1101/2025.03.12.642918

**Authors:** Koshi Maekawa, Nico van Gessel, Ralf Reski, Elena Kozgunova

## Abstract

Although plants share core cell division mechanisms with other eukaryotes, their unique features, such as acentrosomal spindle formation and cytokinesis *via* the phragmoplast, suggest the existence of plant-specific genes. This study used the model bryophyte *Physcomitrium patens*, commonly known as Physcomitrella, to uncover such genes and employed CRISPR/Cas9-based and localization-based screening to identify novel cell division genes. Co-expression data from known mitotic genes were used to create a pool of 216 candidate genes, which were targeted for CRISPR/Cas9 screening. Frameshift mutants with division defects were characterized using high-resolution imaging and fluorophore-based protein localization. Three novel gene families—CYR (Cytokinesis-Related), LACH (Lagging Chromosome), and SpinMi (Spindle and Phragmoplast Midzone)—were identified. CYR is linked to cytokinesis defects, LACH is essential for chromosome segregation, and SpinMi localizes to the spindle and phragmoplast midzone. None of these gene families has homologs in algae, suggesting their emergence during land colonization. These findings demonstrate the utility of co-expression-guided and targeted screening for discovering genes involved in specific cellular processes and provide insights into the evolution of the plant cell division machinery.

## Introduction

Cell division or mitosis involves accurate duplication and distribution of genetic material, followed by the separation of two daughter cells. Errors in cell division are often connected to genetic disorders, cancer, or developmental abnormalities (Gordon *et al*., 2012; McIntosh, 2016). In plants, like in other eukaryotes, cell division is a fundamental process necessary for growth, development, and reproduction. While plants share basic steps of cell division with other eukaryotes, there are also significant differences. For instance, most plant cells lack centrosomes, which serve as microtubule-organizing centres in animals and fungi. As a result, plant spindles are formed without centrosomes, although they remain bipolar (Yamada & Goshima, 2017; Liu & Lee, 2022). Additionally, plant cells are surrounded by rigid cell walls, and cytokinesis occurs through the formation of a cell plate assisted by a cytoskeleton array known as phragmoplast, which transports vesicles containing cell wall materials and promotes cell plate expansion (Smertenko *et al*., 2018; Sinclair *et al*., 2022). Another well-known cytoskeletal array involved in plant cell division is the preprophase band (PPB), which forms before nuclear envelope breakdown (NEBD) and disappears shortly thereafter. This leaves behind a cortical division zone enriched with proteins that guide the phragmoplast during cytokinesis toward the site of cell division (Rasmussen & Bellinger, 2018; Yi & Goshima, 2022; Yamada *et al*., 2025). Notably, in some bryophytes, such as the moss *Physcomitrium patens*, the early development of three-dimensional organs does not rely on the PPB and is presumably governed by different mechanisms (Moody, 2022; Yin *et al*., 2025).

Land colonization by plants was one of the most transformative events in the history of life on Earth. For plants, the transition from water to land brought about dramatic changes in many aspects of biology, including cell division (Horst *et al*., 2016; Chater *et al*., 2016; Reski, 2018a; Resemann *et al*., 2021; Kriegshauser *et al*., 2021; Lüth *et al*., 2023; Knosp *et al*., 2024). It is speculated that ancestors of land plants – related to modern Streptophyte algae – divided through centripetal cleavage, lacked PPB, and their spindles included centrosomes, which also nucleate astral microtubules, a system still observed in modern species like *Klebsormidium* (Buschmann & Zachgo, 2016). The development of unique features of cell division in plants has likely coincided with land colonization and may have been crucial for this event. It has long been suggested that novel genes have emerged in land plants to support the changes in the cell division mechanism (Lipka *et al*., 2015; Rasmussen & Bellinger, 2018; Sablowski & Gutierrez, 2022).

Several plant-specific cell division genes, such as KNOLLE and FASS/TON2, have been identified through forward genetic screening (Traas *et al*., 1995; Lukowitz *et al*., 1996; Camilleri *et al*., 2002; Wright *et al*., 2009). Traditionally, forward genetic mutant collections are generated using methods such as chemical mutagenesis, physical irradiation, or insertional agents like T-DNAs and transposons. However, these approaches are often limited by the need to obtain homozygous progeny and the labour-intensive, time-consuming process of identifying causal mutations responsible for observed phenotypes. The development of CRISPR/Cas9 gene-editing technology has revolutionized targeted mutagenesis, making it more accessible and efficient. CRISPR/Cas9 has also been applied to forward genetic screening in plants (Meng *et al*., 2017; Liu *et al*., 2020).

In this study, we asked if it is possible to identify novel cell division genes in plants by combining CRISPR-based genetic screening with co-expression analysis of known mitotic genes, utilizing the model bryophyte *Physcomitrium patens* (*P. patens*), also known as Physcomitrella (Lueth & Reski, 2023). *P. patens* is a widely used model in plant cell division research owing to its simple body plan, amenability for high-resolution imaging, and ease of genetic manipulation (Reski, 2018b; Rensing *et al*., 2020). Previous studies have functionally characterized numerous cell division genes in *P. patens*, including kinetochore genes (Kozgunova *et al*., 2019) and various microtubule-associated proteins (Nakaoka *et al*., 2012; Miki *et al*., 2014; Kozgunova *et al*., 2022).

Our central hypothesis was that genes involved in a particular cellular process, such as cell division, might share similar expression patterns. Therefore, co-expression data of known cell division genes can be leveraged to enrich a target gene pool with candidate genes likely involved in mitosis. To test this hypothesis, we extracted co-expression information from a publicly available database of 25 known mitotic genes (primarily kinetochore, microtubule-associated and kinase genes), filtered the dataset to focus on genes with unknown functions, and subsequently used CRISPR/Cas9 for genetic screening. From the CRISPR lines, we identified a frameshift mutant with defective phragmoplast formation, resulting in prolonged mitosis and frequent cytokinesis failures. We also discovered a conserved gene family localized to the spindle and phragmoplast midzone and another gene family that is likely essential for moss viability. Knock-down of the latter with inducible RNAi resulted in a chromosome missegregation phenotype. Interestingly, all three gene families are found in land plants, with varying degrees of conservation, but not in the algae lineage. Our findings serve as a proof-of-concept of the effectiveness of targeted gene screens and leveraging co-expression information for gene discovery associated with specific biological processes.

## Results

### A pool of genes for targeted genetic screening

Cell division genes likely make up only a small fraction of the genome. For example, in human cells, it is estimated that approximately 2.4% to 4.25% of genes are involved in cell division (Whitfield *et al*., 2002; Bar-Joseph *et al*., 2008). Given this, we believed that a forward genetic approach is not an efficient strategy to uncover novel cell division genes and decided to pursue a targeted reverse genetic screening instead. By leveraging co-expression data, we aimed to build a pool of previously uncharacterized genes that may be involved in cell division, based on their co-expression with known mitotic genes (Figure 1). Multiple mitotic genes, including kinetochore genes (Kozgunova, 2025), microtubule-associated proteins (Nakaoka *et al*., 2012; Kosetsu *et al*., 2013; Leong *et al*., 2018; Kozgunova *et al*., 2022) and motor proteins (Miki *et al*., 2014; Wu & Bezanilla, 2014) have been functionally characterized in Physcomitrella. We used the publicly available JGI Gene Atlas database to collect information on co-expression networks of selected mitotic genes (**Supplemental Table 1**). As an arbitrary cut-off score, we adopted a co-expression value above 0.8. To further narrow down our gene pool, we excluded genes whose function could be predicted based on homology and genes that had been functionally characterized based on publication records. In addition, we selected several Arabidopsis genes that have been reported to be highly expressed in dividing cells, either based on synchronized culture (Menges *et al*., 2002) or embryo analysis (Slane *et al*., 2014), but were not further characterized. We then identified homologs of these genes in Physcomitrella and included them in the screening. In total, 52 genes were selected based on the Arabidopsis studies, while the remaining 164 genes were identified through co-expression analysis. A full list of the in total 216 genes selected for screening, along with the mitotic genes they co-express with, is provided in **Supplemental Table 1**.

**Figure 1.**
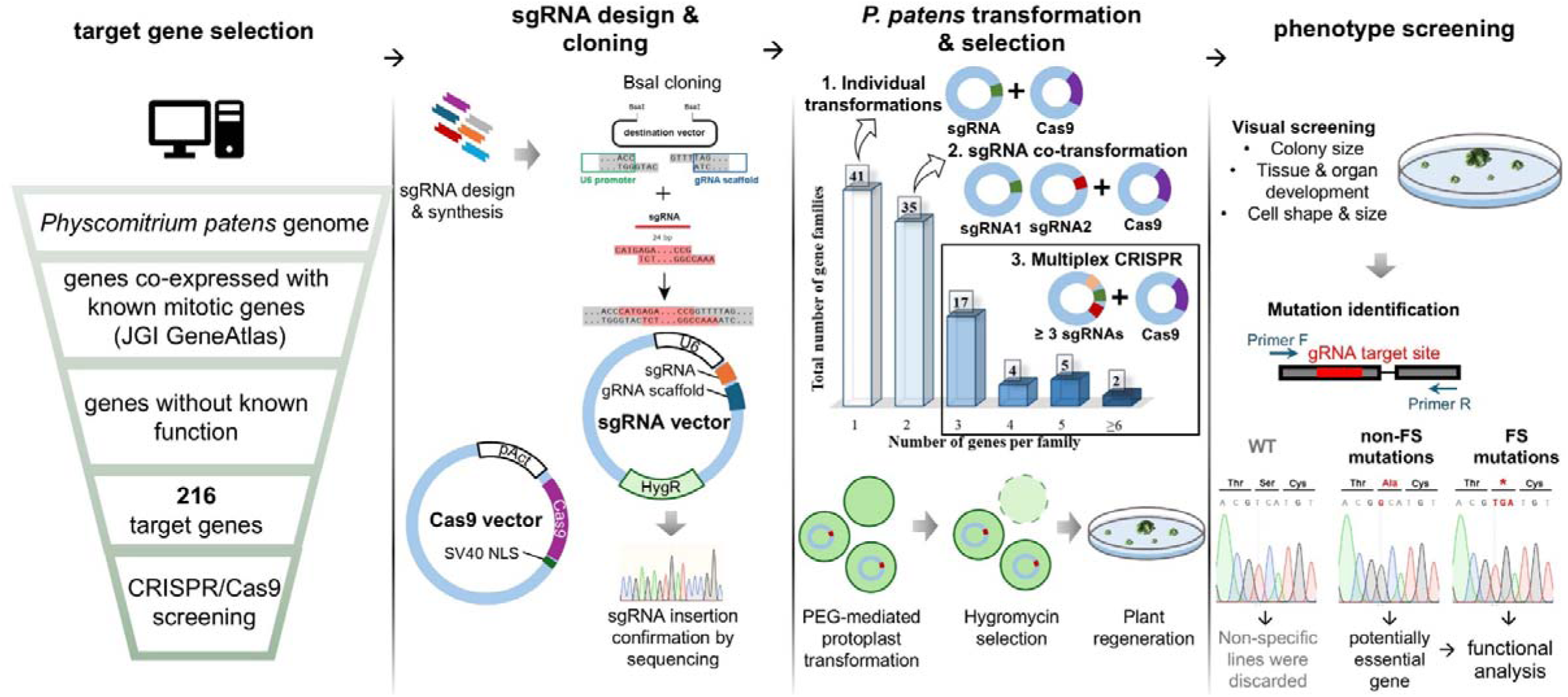
Experimental design: gene selection and CRISPR screening workflow.

### A gRNA library and screening

Guide RNAs (gRNAs) were designed manually using the CRISPOR website (Concordet & Haeussler, 2018), targeting exon regions, typically choosing the exon that was present in all transcript variants (**Supplemental Table 1**). In addition, the genes were grouped based on their homology, and when possible, a single gRNA was designed to target multiple genes from a single gene family through a conserved site. gRNA expression was driven by the U6 promoter and gRNA integration site was followed by gRNA scaffold (Lopez-Obando *et al*., 2016) (**Figure 1**). When three or more gRNAs were required to target all identified gene homologues, we performed an additional cloning step to incorporate all gRNAs into a single vector. As a background line, we chose a line expressing GFP-tubulin and H2B-RFP, which allows us to visualise microtubules and chromosomes, respectively (Nakaoka *et al*., 2012). Hereinafter, we refer to this line as the GH line.

### Phenotype screening and mutation mapping

After transforming the gRNA vectors into moss protoplasts, we visually assessed the phenotypes of the regenerated colonies that survived antibiotic selection. We hypothesized that defects in cell division would result in smaller colonies and abnormal cell or organ shapes. Accordingly, we selected colonies exhibiting these phenotypic characteristics and sequenced them.

Upon sequencing, we have encountered three potential outcomes:

- No mutation was detected at the gRNA target site, which was considered a non-specific result, leading to the exclusion of such colonies.
- Exclusively non-frameshift mutations were identified in multiple colonies. This outcome suggested that the targeted gene might be essential for moss viability, and in some cases, we pursued further analysis of such genes using alternative methods.
- Frameshift mutations in the gRNA target site were correlating with the observed phenotype, providing strong evidence of a functional relationship between the mutation and the phenotype.

One of the gene families was shown to have severe growth retardation after frameshift mutation (**Figure 2 A, B**). We decided to further pursue functional characterization of this gene family.

**Figure 2.**
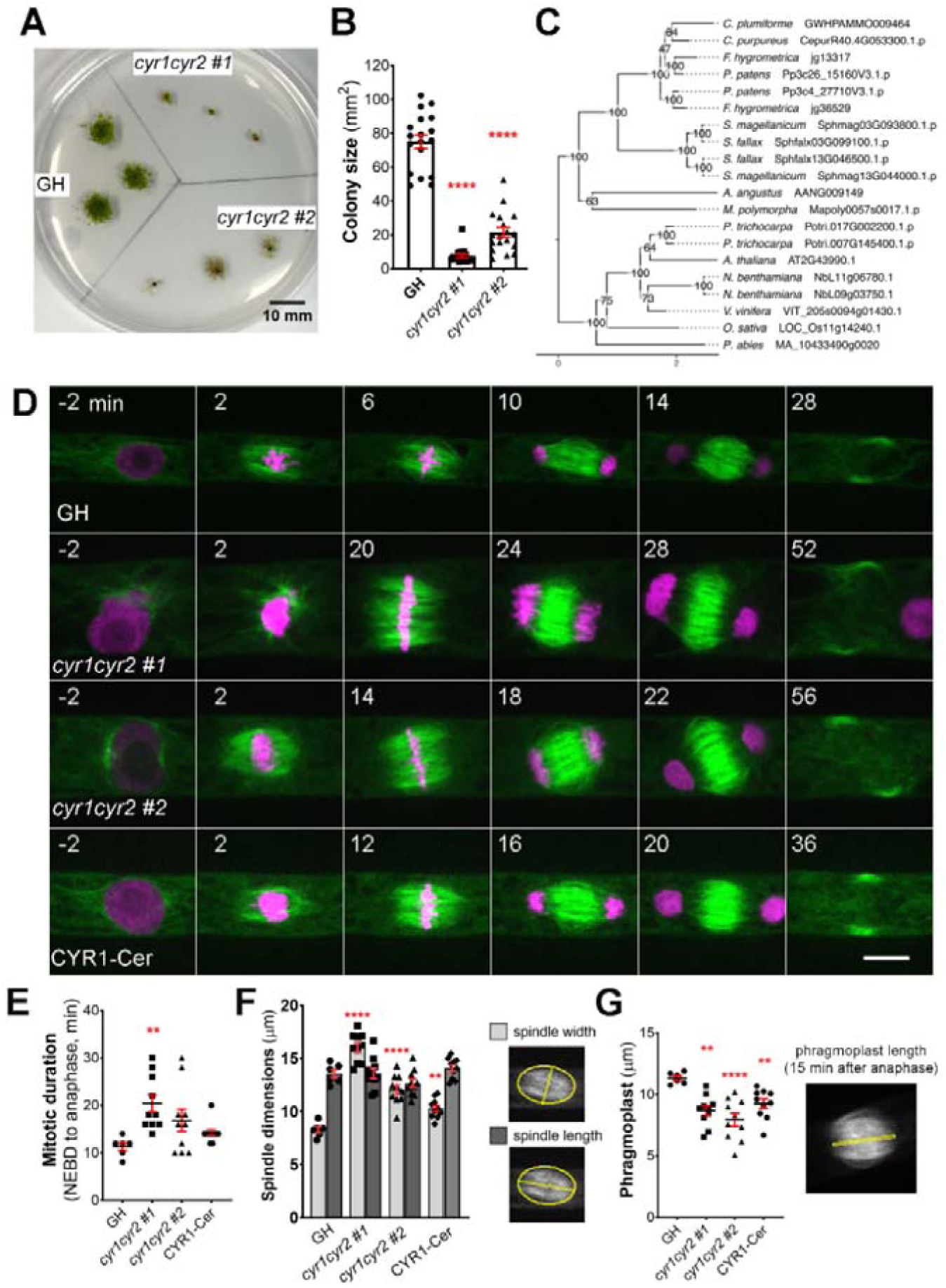
Frameshift mutations in the cytokinesis-related gene family produce various abnormalities during cell division in protonema cells in *P. patens*. (**A**) Representative images of moss colonies after 4 weeks of culture: GH is a control, *cyr1cyr2 #1* and *#2* are independent, adapted frameshift lines (phenotypes weakened over time) isolated through CRISPR genetic screening. Scale, 10 mm. (**B**) Colony size measured after 4 weeks of culture. Each data point corresponds to a single colony (mean□±□SEM, ****p≤□0.0001 by one-way ANOVA with Dunnett’s multiple comparison test against GH) (**C**) Phylogenetic reconstruction of the *CYR* gene family based on maximum-likelihood tree inference with midpoint-rooting. Values at internal nodes indicate percentage support by 1000 bootstrap samples. (**D**) Representative images of cell division in GH, *cyr1cyr2 #1*, *cyr1cyr2#2* and CYR1-Cerulean (rescue line in the *cyr1cyr2#2* background). Time zero is set at NEBD, and single focal plane images are shown. Scale, 10 µm. (**E**) Mitotic duration calculated from NEBD to anaphase onset. Each data point corresponds to a single cell (mean□±□SEM, \*\**p*□=□0.041 by one-way ANOVA with Dunnett’s multiple comparison test against GH) (**F**) Spindle dimensions: spindle width (light gray bars) and spindle length (dark gray bars). Each data point corresponds to a single cell (mean□±□SEM, ****p≤□0.0001, \*\**p*□=□0.064 by one way ANOVA with Dunnett’s multiple comparison test against GH). No statistically significant difference in spindle length found. (**G**) Phragmoplast length in GH, *cyr1cyr2 #1*, *cyr1cyr2#2* and CYR1-Cerulean (rescue line in the *cyr1cyr2#2* background). Each data point corresponds to a single cell (mean□±□SEM, \*\**p*□=□0.010, ****p≤□0.0001, \*\**p*□=□0.095 by one-way ANOVA with Dunnett’s multiple comparison test against GH).

### Cytokinesis-related (CYR) gene family affects spindle and phragmoplast

This gene family consists of two genes in *P. patens* and potential homologues were also found in other bryophytes, as well as in some vascular plant species. We named this gene family <u>Cy</u>tokinesis-<u>R</u>elated (*CYR*) to reflect the cellular phenotype observed in frameshift mutants, with *CYR1* (version 3.3: Pp3c26_15160; version 6.1: Pp6c26_8300) and *CYR2* (version 3.3: Pp3c4_27710; version 6.1: Pp6c4_14480) assigned as the respective gene names. The two genes in this family encode proteins of approximately 100 kDa, with around 57% shared amino acid sequence identity and an assignment to the PANTHER family GPI-ANCHORED ADHESIN-LIKE PROTEIN (PTHR36022). Using CYR1 and CYR2 as queries we performed homology searches against a reference species set covering major lineages of Archaeplastida, including chlorophytic and charophytic algae. Careful evaluation and phylogenetic reconstruction revealed conservation of the gene family across land plants (**Figure 2 C**), but no homologs in algae. *In silico* prediction of subcellular localization for both protein sequences identified no specific signal peptides, but hinted towards cytoplasmic and nuclear localization (**Supplemental Table 2**). In our co-expression analysis, the CYR gene family was selected based on co-expression with mitotic kinase MPS1 (Pp3c6_320) and microtubule-associated protein TPX2-4 (Pp3c23_4540) (**Supplemental Table 1**). To the best of our knowledge, this gene family has not been characterized previously.

During CRISPR screening we isolated two independent clones in which both *CYR1* and *CYR2* genes carried frameshift mutations (**Supplemental Figure 1A**). To assess their cellular phenotypes, we first stained the cells with FM4-64, a membrane marker, and observed incomplete cell plate formation and multinucleated cells, suggesting cytokinesis failure (**Supplemental Figure 1B**). High-resolution imaging of *cyr1cyr2* mutants confirmed cytokinesis failure in 2 out of 3 cells in *cyr1cyr2 #1* line and 2 out of 4 cells in *cyr1cyr2 #2* line (**Supplemental Video 1**). However, due to the severity of the phenotype, it was difficult to collect data from a large number of cells.

After continuous culture, the phenotype became less pronounced, possibly due to adaptive mutations, epigenetic changes, or polyploidization. In these “adapted” lines, cytokinesis failure was no longer observed during live-cell imaging. To further characterize their morphology, we repeated FM4-64 staining and quantified the frequency of abnormal phenotypes, including multinucleated cells, incomplete cell plates, multiple cell plates, and abnormal branching (**Supplemental Figure 1C–D**). Overall, approximately 32% of cells in the adapted *cyr1cyr2* #1 line and 47% in the adapted *cyr1cyr2* #2 line exhibited some form of abnormality. However, multinucleated cells, which typically represent complete cytokinesis failure, accounted for only 6.7% and 14.8% of cells in the two lines, respectively, suggesting that complete cytokinesis failure was relatively uncommon. Nonetheless, we proceeded with quantitative analyses using high-resolution live-cell imaging on these “adapted” lines. Both *cyr1cyr2 #1* and *#2* lines showed a delay in mitotic progression compared to the control (**Figure 2E**). Although frameshift mutants could form a bipolar spindle, spindle proportions were distorted. The average spindle width in *cyr1cyr2 #1* and *cyr1cyr2 #2* frameshift lines was 16.03 µm±0.43 and 12.07 µm±0.43, respectively, while in control GH line average spindle width was 8.28 µm±0.27. On the other hand, spindle length did not change significantly (GH 13.57 µm±0.37, *cyr1cyr2 #1* 13.60 µm±0.47, *cyr1cyr2 #2* 12.66 µm± 0.38) (**Figure 2F**). Phragmoplasts were narrower in frameshift mutants at 8.66 µm±0.39in *cyr1cyr2 #1* and 7.92 µm±0.53, in *cyr1cyr2 #2,* compared to the control 11.32 µm± 0.18 (**Figure 2G**). To verify the phenotype of the frameshift mutants, we ectopically expressed CYR1-Cerulean under the pEF1α promoter (Aoyama *et al*., 2012) in the *cyr1cyr2 #2* background. Although the phenotype was not completely restored to wild type, CYR1-Cerulean showed a trend toward rescue in our quantifications (**Figure 2 D-G**). For example, the average spindle width in the CYR1-Cerulean line was reduced to 10.28 µm±0.29 and the average phragmoplast increased to 9.22 µm±0.38. We speculate that the incomplete rescue might be due to differences in expression levels between the original gene and the ectopically expressed rescue construct. It is also possible that, since the frameshift line has evolved over time, other adaptive mechanisms may affect cell division phenotypes.

We have also occasionally observed abnormal phragmoplast formation, such as a disorganized phragmoplast or multiple phragmoplasts forming in the “adapted” frameshift lines. This phenotype was observed in 2 out of 11 cells in *cyr1cyr2 #1* and in 4 out of 12 cells in the *cyr1cyr2 #2* line (**Supplemental Video 2**).

To further assess the potential redundancy between *CYR1* and *CYR2*, we also examined the phenotype of single frameshift mutants, generated using the same gRNAs (**Supplementary Figure 2A**). In contrast to the double mutants, neither *cyr1* nor *cyr2* single mutants displayed strong growth defects, although quantitative analysis showed that *cyr 1 #30* colonies were slightly smaller comparing to GH (**Supplementary Figure 2B, C**). Closer investigation of cell division by live-cell imaging did not find any abnormalities in *cyr1* and *cyr2* single mutants (**Supplementary Figures D–G**). These findings support the interpretation that CYR1 and CYR2 have redundant roles in cell division.

We next aimed to visualize the localization of CYR1 and CYR2 in living cells of the wild-type. In *P. patens*, the high rate of homologous recombination enables the creation of endogenously tagged lines, where the tagged protein replaces the native protein (Mueller *et al*., 2014; Lüth *et al*., 2023). To visualize the cellular localization of the CYR proteins, we attempted endogenous gene tagging using the mScarlet fluorophore. Although we successfully isolated and verified the tagged lines through genotyping PCR, we were unable to detect fluorescent signals, even with high laser power and extended exposure times. This suggests that CYR proteins are weakly expressed under normal conditions. Next, we constructed vectors for ectopic expression of CYR1 or CYR2 fused to Citrine under the control of the rice actin promoter, which enabled us to observe the localization of these proteins. Both CYR1-Citrine and CYR2-Citrine exhibited cytoplasmic localization prior to nuclear envelope breakdown (NEBD). During cytokinesis, both CYR1 and CYR2 showed bright signals at the phragmoplast midzone and weaker signals throughout the rest of the phragmoplast. These signals expanded as the phragmoplast grew, ultimately fusing with the cell boundaries (**Fig 3A, B; Supplemental Video 3**). Interestingly, we also occasionally observed multiple phragmoplast formation in the CYR1-Citrine (*n* = 2 out of 7), but not in the CYR2-Citrine overexpression line (*n* = 7) (**Supplemental Video 2**). While the mechanism remains unclear, no cytokinesis failure or growth defects were detected in these lines.

**Figure 3.**
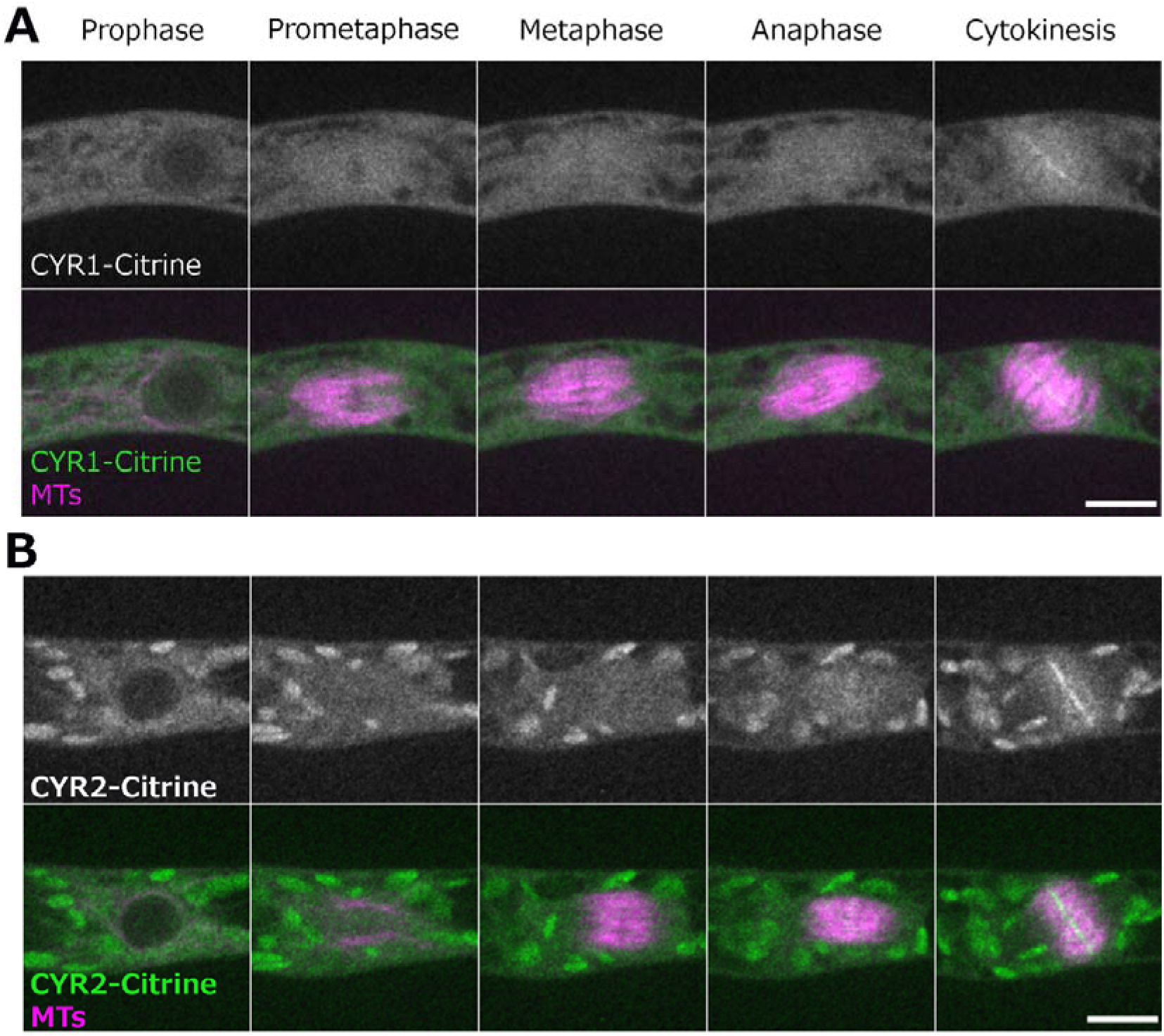
Localization of CYR1 and CYR2 during cell division. Live imaging of wild-type *P. patens* caulonemal apical cells expressing mCherry-tubulin (magenta) and (**A**) CYR1-Citrine (green) or (**B**) CYR2-Citrine (green). CYR1-Citrine and CYR2-Citrine are ectopically expressed under a rice actin promoter. Images were acquired every 2 min as a Z-stack (5 µm in 2.5 µm steps); best focal plane is shown. Scale, 10 µm.

Taken together, the combination of the cellular phenotype observed after frameshift mutations and the cellular localization data suggests that the CYR gene family function is related to cytokinesis in Physcomitrella. However, the specific role of these genes and their interacting partners remain to be identified.

### Lagging Chromosome (LACH) gene family is essential

Another gene family, named <u>La</u>gging <u>Ch</u>romosome (*LACH*) after the later discovered cellular phenotype, was originally recognized in our CRISPR screening as an essential gene candidate. During CRISPR screening we were able to isolate several non-frameshift mutants, but not plants containing frameshift mutations, suggesting that frameshift mutations in the *LACH* gene family are lethal in *P. patens*. We found the *LACH* gene to be co-expressed with mitotic kinases Haspin (Pp3c5_21110) and MPS1 (Pp3c6_320) (**Supplemental Table 1**). In addition to the *LACH* gene in Physcomitrella (version 3.3: Pp3c25_4280; version 6.1: Pp6c25_7230), phylogenetic reconstruction within our species set revealed potential homologs of this gene in other bryophytes, the clubmoss *Diphasiastrum complanatum* and the gymnosperm *Picea abies* (**Figure 4A**). Protein functionality could not be clearly predicted and no family assignment was inferred by alignment against the PANTHER database. Our *in-silico* analysis revealed a high probability for localization to the cytoplasm and nucleus (**Supplemental Table 2**). Since we could not isolate a frameshift line using CRISPR/Cas9, we decided to pursue further characterization of this gene using different methods. To investigate the localization of LACH protein in living cells, we generated an endogenously tagged line in which the *LACH* gene was fused to the mNeonGreen fluorophore. LACH-mNeonGreen signals were observed in the cytoplasm and nucleus during interphase. After NEBD, we observed accumulation of brighter signals in the centre of the spindle during prometaphase and metaphase, although overall fluorescent signals intensity was low. These signals disappeared upon the anaphase onset **(Figure 4B, Supplemental Figure 3**).

**Figure 4.**
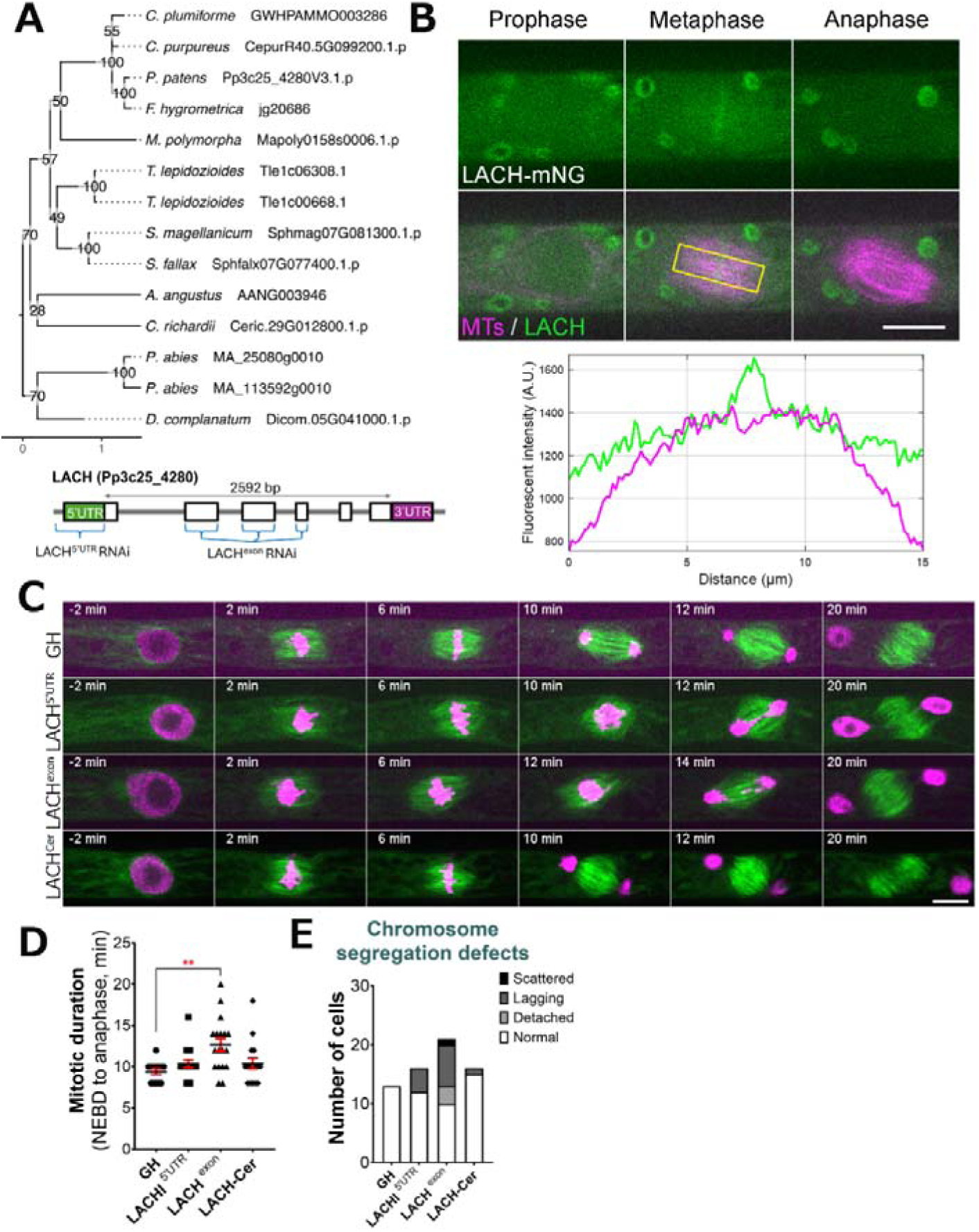
LACH localization and phenotype after RNAi knock-down. (**A**) (*top*) Phylogenetic reconstruction of the LACH family based on maximum-likelihood tree inference with midpoint-rooting. Values at internal nodes indicate percentage support by 1000 bootstrap samples. (*bottom*) Gene model of the LACH gene and sites targeted by RNAi. (**B**) Live imaging of *P. patens* protonemal apical cells expressing mCherry-tubulin (magenta) and LACH-mNeonGreen (green). Plot shows fluorescent intensity distribution (A.U.) in metaphase cell; measured area is indicated with a yellow rectangle. Full localization data can be found in Supplemental Figure 3. Scale, 10 µm. (B) Representative mitotic progression and chromosome missegregation in the LACH RNAi lines. ‘GH’ is the control line, LACH^Cer^ is a rescue line expressing codon-replaced LACH-Cerulean in the LACH^exon^ RNAi background. Scale, 10 µm. (**D**) Mitotic duration calculated from NEBD to anaphase onset. Each data point corresponds to a single cell (mean□±□SEM, \*\*\**p*□=□0.0008 by one way ANOVA with Dunnett’s multiple comparison test against GH) (**E**) Frequency of chromosome missegregation defects in LACH RNAi lines. Chromosome missegregation defects were classified into three types: chromosomes detached from the metaphase plate (detached chromosomes), lagging chromosomes in anaphase (lagging chromosomes), and scattered (severe chromosome missegregation). Y-axis shows the actual number of cells observed.

Additionally, we aimed to functionally characterize the *LACH* gene using an inducible RNA interference (RNAi) system, in which RNAi production is triggered by the addition of β-estradiol to the culture medium (Kubo *et al*., 2013). Since the gene knock-down is triggered only in the presence of β-estradiol, this system is widely used to study essential genes in Physcomitrella (Nakaoka *et al*., 2012). We designed two independent RNAi constructs: one targeting the 5’ UTR region and another targeting an exon region of the *LACH* gene. After line selection, we observed phenotypic changes following 5-6 days of culture with 1 µM β-estradiol in the growth medium, because prolonged culture can complicate data interpretation due to side effects. Overall, we did not notice changes in cell growth or cell shape comparing to the control. A closer investigation of cell division showed that both RNAi constructs led to various chromosome missegregation phenotypes, including detached and/or lagging chromosomes during mitosis (**Figure 4C, E**). The chromosome missegregation phenotype was more pronounced in the exon RNAi line (**Figure 4C, Supplemental Video 4**) and similarly, mitotic delay was also more noticeable in the LACH exon RNAi line, which we attribute to the strength of the RNAi line and efficiency of gene knock-down (**Fig 4D, E**). To confirm the specificity of this phenotype, we generated a rescue line expressing codon-replaced LACH-Cerulean in the background of the exon RNAi construct. This transgene restored normal mitotic timing and significantly reduced the lagging chromosome phenotype, supporting that these defects are due to *LACH* depletion (**Figure 4C–E**).

Based on the RNAi phenotype and its localization, we conclude that *LACH* function is related to regulating chromosome segregation during mitosis in *P. patens*.

### SpinMi gene family, identified through localization screening

Besides CRISPR-based genetic screening, we chose 21 gene candidates for additional localization screening. Candidates were selected based on their co-expression with multiple known mitotic genes during the initial gene search (**Supplemental Table 1**). We speculated that their repeated appearance in our co-expression analysis suggests a higher likelihood that they are involved in cell division.

For the localization screening, we created endogenously tagged versions of these genes by fusing them with the mNeonGreen fluorophore in a background line expressing the microtubule marker mCherry-α-tubulin (Kosetsu *et al*., 2017). After confirming the correct integration through genotyping PCR, we performed live-cell imaging in protonema cells to observe protein localization. One of the tagged proteins was distinctly observed at the mitotic spindle and phragmoplast during cell division, prompting us to conduct further analysis. Based on the protein localization, we designated this gene family as *SpinMi* (<u>Spin</u>dle and Phragmoplast <u>Mi</u>dzone).

The *SpinMi* gene family contains a single-copy gene in Physcomitrella (version 3.3: Pp3c14_17140; version 6.1: Pp6c14_9440) and appears to be highly conserved across land plants, including vascular plants, with species-specific expansions (**Figure 5A**). However, similar to the *CYR* and *LACH* gene families, we were unable to detect clear homologs in algae. All identified homologs share an assignment to the PANTHER family OS03G0691500 PROTEIN (PTHR45287) and our *in-silico* predictions revealed hints for localization to the cytoplasm and nucleus (**Supplemental Table 2**).

**Figure 5.**
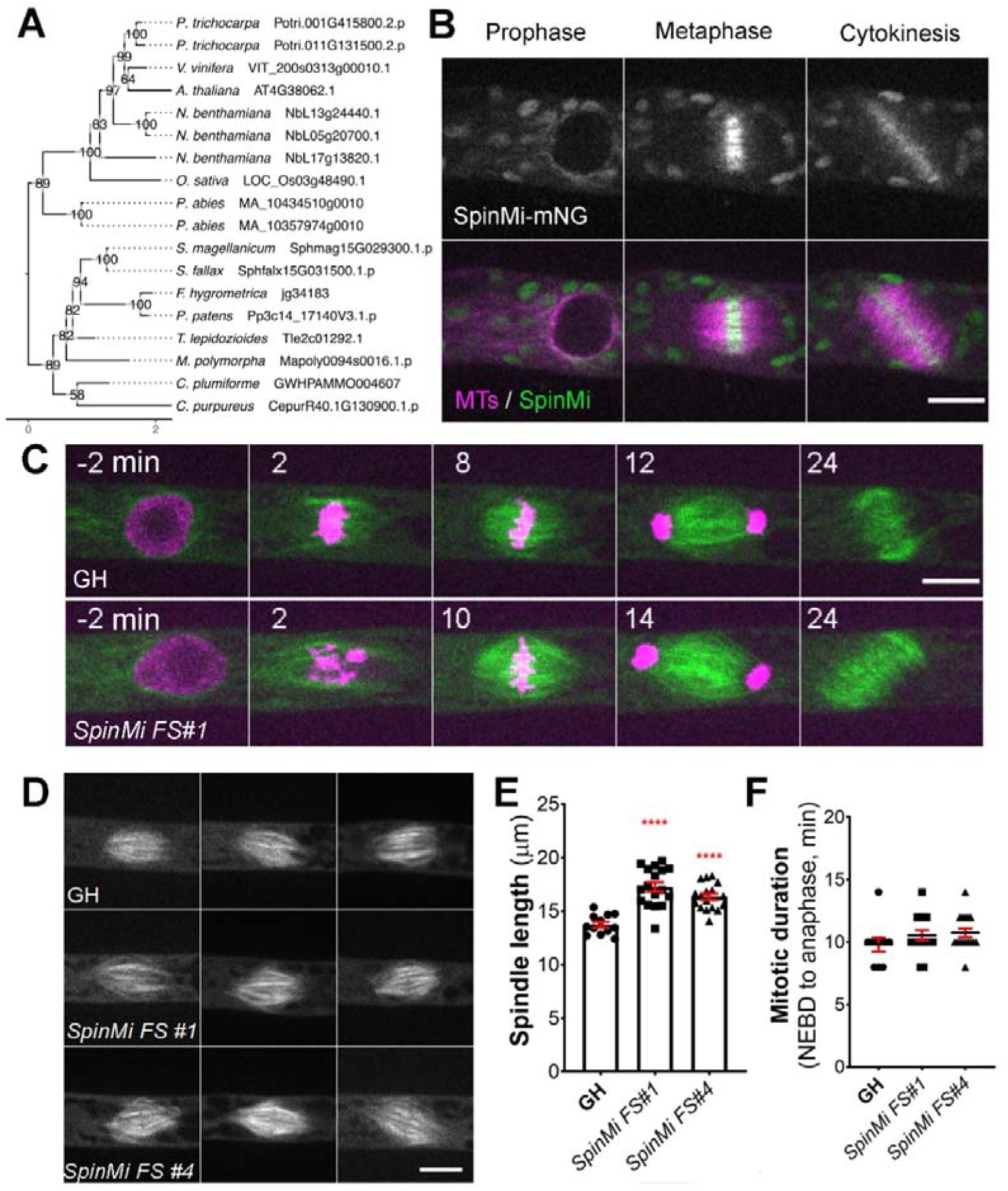
SpinMi localization and frameshift phenotype. (**A**) Phylogenetic reconstruction of the SpinMi family based on maximum-likelihood tree inference with midpoint-rooting. Values at internal nodes indicate percentage support by 1000 bootstrap samples. (**B**) Live imaging of *P. patens* protonemal apical cells expressing mCherry-tubulin (magenta) and SpinMi-mNeonGreen (green). Full localization data can be found in Supplemental Figure 3. Scale, 10 µm. (**C**) Representative images of cell division in GH and *SpinMi FS#1* (frameshift mutant line). Scale, 10 µm. (**D**) Representative images of metaphase spindles in control (GH), *SpinMi FS#1* and *SpinMi FS#2* mutant lines. Three spindles per each line are shown. Scale, 10 µm. (**E**) Metaphase spindle length measured in GH, *SpinMi FS#1* and *SpinMi FS#2*. Each data point corresponds to a single cell (meanD±DSEM, \*\*\*\**p*D≤D0.0001 by one way ANOVA with Dunnett’s multiple comparison test against GH) (**F**) Mitotic duration from NEBD to anaphase onset in GH, *SpinMi FS#1* and *SpinMi FS#2* lines. No statistically significant difference found. Each data point corresponds to a single cell (mean ± SEM).

During prophase, SpinMi-mNeonGreen was distributed throughout the cytoplasm and was also observed on microtubule bundles that surround the nucleus before NEBD in protonema apical cells (Yin *et al*., 2025). Following NEBD, SpinMi-mNeonGreen was observed at the forming mitotic spindle, with a higher fluorescent intensity broadly localized at the spindle midzone. This localization persisted until anaphase, and afterwards SpinMi-mNeonGreen relocated to the forming phragmoplast, with brighter fluorescence observed in the phragmoplast midzone (**Figure 5B, Supplemental Figure 4, Supplemental Video 5**). To assess whether the localization of SpinMi during mitosis is microtubule-dependent, we treated dividing cells with 20 µM oryzalin, a microtubule-depolymerizing drug (Pressel *et al*., 2008). Following oryzalin addition, SpinMi-mNeonGreen signals gradually dissipated following microtubule depolymerization, indicating that SpinMi localization depends on microtubules during cell division (**Supplemental Figure 4C**).

Next, we sought to investigate the function of SpinMi during mitosis. We isolated two independent frameshift mutants using CRISPR/Cas9 (**Supplemental Figure 5A**). Upon initial observation, the frameshift mutants appeared to grow normally without any visible defects compared to controls (**Supplemental Figure 5 B, C**). This potentially explains why the *SpinMi* gene was not identified in our CRISPR screening, which focused on targeting moss colonies with noticeable growth defects.

We next examined the cellular phenotype of *spinmi* frameshift mutants during mitosis in protonema cells. Mitotic duration, chromosome segregation, phragmoplast assembly, and cytokinesis all appeared normal in the frameshift mutants (**Figure 5 C, F**). However, upon closer inspection, we observed that the mitotic spindles in *spinmi* frameshift mutants were slightly elongated compared to the control (*SpinMi FS#1* 17.27 µm±0.43, *SpinMi FS#2* 16.24 µm±0.29 vs GH 13.74 µm±0.26) (**Figure 5 D, E**). Despite this, no detrimental effects on cell division were evident under normal conditions. *SpinMi* may contribute to fine-tuning spindle architecture, as suggested by its localization and the observed subtle increase in spindle length in mutant lines.

## Discussion

This study was originally designed as a proof of concept that genetic screening combined with the analysis of co-expression data has the power to identify novel genes associated with specific cellular process, specifically cell division, in the model bryophyte *P. patens*. In the context of cell division, expression profiles from synchronized cell cultures in both animal and plant models are often employed to determine which genes are involved in this process (Menges & Murray, 2002; Bar-Joseph *et al*., 2008). While this method has its merits, it also presents challenges; for instance, not all plant models have established protocols for cell cycle synchronization, and such treatments might be disruptive, potentially affecting gene expression.

We utilized publicly available co-expression data from the Phytozome database, specifically the JGI Plant Gene Atlas project, which contains comprehensive transcriptome data from various growth conditions, different developmental stages (protonema, gametophore, sporophyte, etc.), and multiple laboratories (Perroud *et al*., 2018; Sreedasyam *et al*., 2023). We hypothesized that even without cell synchronization, cell division genes might exhibit co-expression under similar conditions and/or life stages. Among the genes we selected for screening, we identified three novel gene families that are likely associated with cell division in the moss *P. patens*. To the best of our knowledge, these gene families have not been functionally characterised previously and homology inference did not provide sufficient insights. However, phylogenetic analysis allowed us to trace the ancestry of all three gene families back to early land plants, including liverworts and the moss *Takakia*, which is sister to all other mosses (Hu *et al*., 2023). Strikingly, we did not find clear homologs in Streptophyte algae, the closest relatives to land plants (Hess *et al*., 2022). This raises the question of whether the emergence of the families described in our study is a consequence of the transition to land colonization. Furthermore, the functional roles of these genes remain enigmatic, as predicted functionalities of two of them (*CYR* and *LACH*) do not match any well-characterized PANTHER annotations and no functional domain annotation was found. This suggests that their cellular function may involve yet-to-be-understood mechanisms.

Our findings regarding the *CYR* gene family indicate a strong association with cytokinesis in *P. patens*. The cellular localization of CYR1 and CYR2 at the phragmoplast midzone, along with the cytokinesis defects observed in frameshift mutants, suggests that these proteins play a role in phragmoplast microtubule dynamics and/or cell plate formation. While their precise molecular function remains unknown, their localization pattern is reminiscent of secretory carrier membrane proteins (SCAMPs) and kinesin12-II, which drives vesicle transport in the *P. patens* phragmoplast overlapping anti-parallel microtubules in the phragmoplast (Yamada *et al*., 2025). Unlike kinesin12-II, CYR proteins do not contain known microtubule-binding domains or motor motifs, its annotation as GPI-anchored protein suggests that it may be a membrane-bound protein.

Interestingly, we occasionally observed multiple phragmoplast formations in the CYR1-Citrine overexpression line and *cyr1cyr2* frameshift mutants, but not in the CYR2-Citrine line (**Supplemental Video 2**). It is possible that either overexpression or fluorophore tagging of CYR1, or a combination of these factors, acts in a dominant-negative way and can recapitulate some of the cellular phenotypes observed in the CRISPR frameshift mutants. However, we did not observe cytokinesis failure or growth defects, suggesting that negative effects remain mild.

The incomplete rescue observed in ectopic CYR1-Cerulean expression lines suggests that tight regulation of CYR protein expression levels may be necessary for proper function. Further biochemical and genetic studies will be required to clarify their precise molecular roles.

Our characterization of the *LACH* gene family suggests a function related to chromosome segregation. CRISPR-induced mutants for LACH contained exclusively in-frame mutations. We did not attempt homology-directed knock-in of a stop codon, but our inability to recover null mutants, combined with RNAi phenotypes, supports a role for LACH in essential mitotic functions. The fact that we were unable to isolate viable *LACH* frameshift mutants further supports the notion that *LACH* is crucial for the moss, and that its loss is lethal. The localization of LACH protein at the spindle midzone and its loss-of-function phenotype, which includes chromosome missegregation and mitotic delay, strongly suggest an involvement in chromosome cohesion and/or microtubule-kinetochore attachment. During cell division, kinetochore proteins show clear dot-like localization patterns in Physcomitrella (Kozgunova *et al*., 2019), which was not observed in the LACH-mNeonGreen line. Therefore, we suggest that the LACH protein is not a part of the kinetochore complex. However, it is possible that LACH is indirectly involved in centromere/kinetochore formation. For example, in Arabidopsis cenH3 histone chaperone NASP does not show a clear centromeric localization, but its reduced expression affects deposition of centromeric histone H3 (Le Goff *et al*., 2020). Another possibility is that LACH function is related to the cohesin complex, which mediates cohesion between replicated sister chromatids and is essential for chromosome segregation in dividing cells. Cohesin complex role during mitosis have not been investigated in Physcomitrella, however in maize, cohesin subunit Structural Maintenance of Chromosome3 (SMC3) mutants show premature loss of sister chromatid cohesion and chromosomes missegregation (Zhang *et al*., 2020a). Future research could explore whether *LACH* functions through direct interactions with cohesion regulators or kinetochores.

The third gene family, *SpinMi*, was not identified through CRISPR screening, but rather through localization analysis. This highlights a limitation of our CRISPR screening approach: genes that do not exhibit a strong phenotype after mutation are less likely to be identified using our workflow, which focuses on colonies with growth defects or candidates for essential genes (as indicated by the absence of frameshift mutations). Conducting a localizome screening can serve as either an alternative to, or a complement for, CRISPR screening. In *P. patens*, SpinMi-mNeonGreen localized to the spindle and phragmoplast midzone in a microtubule-dependent manner. Microtubule polarity in mitosis is typically oriented with most of microtubules plus-ends directed toward the spindle and phragmoplast midzone (Dhonukshe *et al*., 2006). Therefore, it is possible that SpinMi preferentially binds to the plus-ends of microtubules or forms a complex with other plus-end-binding proteins. Although *spinmi* frameshift mutants did not exhibit severe defects in mitotic progression, the subtle increase in spindle length suggests that SpinMi plays a role in fine-tuning microtubule organization during mitosis. For example, elongated spindles have also been observed following the knockdown of microtubule-nucleating proteins such as γ-tubulin and augmin in Physcomitrella and Arabidopsis (Nakaoka *et al*., 2012; Romeiro Motta *et al*., 2024). The loss of *SpinMi* may alter the dynamic properties of microtubules, potentially affecting the balance of forces exerted by motor proteins and microtubule-associated proteins that regulate spindle assembly. Given that *SpinMi* homologs are highly conserved in land plants, it is also of interest to investigate their functions in other model plants and in meiosis.

Our findings offer insights into the genetic components of cell division in early land plants and provide a workflow that integrates CRISPR-based genetic screening with co-expression analysis for gene discovery in the popular model bryophyte *P. patens*. Compared to traditional forward genetics screening, our approach bypasses the need for large-scale mutagenesis and segregation analysis by targeting high-confidence candidates directly. Although transformation and genotyping are still required, the co-expression-based filtering dramatically reduces the search space. Nonetheless, we acknowledge that genes producing weak or lethal phenotypes are likely to be missed in our workflow.

The identification of three novel gene families, unique to land plants, suggests that their emergence was linked to the evolutionary transition from aquatic to terrestrial environments. This transition likely required significant adaptations to the cell division machinery, in response to the dramatic changes in the environment and the need for complex organ development. Moving forward, future studies can focus on further functional analysis of *CYR*, *LACH* and *SpinMi* gene families, investigating their potential interactions with known mitotic regulators, and exploring their functional conservation across different plant lineages to better understand how they contribute to plant cell division and growth.

## Materials and Methods

### *P. patens* culture and transformation

We used *P. patens* lines derived from the Gransden strain (International Moss Stock Center IMSC, accession number 40001), which were maintained at 25 °C under continuous light conditions. Protoplast transformation was carried out according to Hohe *et al*. (2004) using protoplasts prepared from protonema tissue after 6–7 days of culture. To isolate protoplasts, the protonemata was incubated in a solution containing 2% driselase and 8% (w/v) mannitol for 30 min at room temperature with gentle rocking. The resulting suspension was filtered through a 50 µm nylon mesh and centrifuged at 180 × g for 2 min to pellet the protoplasts. After discarding the supernatant, the protoplasts were washed twice with 20 mL of 8% mannitol. Prior to transformation, the protoplast concentration was adjusted to 1.6□×□10□ cells per mL using MMM solution (0.1% MES, pH 5.6, 15 mM MgCl□, and 9.1% (w/v) mannitol). Next, 300□µL of the prepared protoplasts were combined with 30□µL of plasmid DNA (30□µg) and 300□µL of 40% (w/v) polyethylene glycol solution (10□mM Tris-HCl, pH 8.0, 100□mM Ca(NO□)□, and 8% mannitol). For gRNA transformation, equal amounts of circular gRNA plasmid and *Streptococcus pyogenes* Cas9 expression vector were simultaneously transferred into the moss. The mixture was incubated at 45□°C for 5 min, followed by 10 min at 20□°C, and subsequently maintained overnight at 25□°C in darkness. The transformed protoplasts were then plated on protoplast regeneration medium plates overlaid with sterile cellophane. After three days of growth, the regenerated protoplasts were transferred onto BCDAT medium containing the appropriate antibiotics for selection (Hohe *et al*., 2004; Yamada *et al*., 2016). Genomic integration of transgenes was verified through genotyping PCR and/or sequencing for CRISPR-edited lines. For transformation experiments, the GH line, which expresses histone H2B-mRFP and GFP-α-tubulin, was utilized for CRISPR and RNAi studies (Nakaoka *et al*., 2012). The line expressing mCherry-α-tubulin under *EF1_α_* promoter was used as a background line for Citrine and mNeonGreen tagging experiments (Kosetsu *et al*., 2017). A complete list of the transgenic lines generated in this study is provided in **Supplemental Table 3**.

For colony growth assays, protoplasts were prepared from moss protonemata using driselase treatment and mannitol wash, following the same protocol as used for gene transformation. Following 2–3 weeks of protoplast regeneration, individual colonies were selected and transferred to standard BCDAT agar plates for further growth. Since each colony is presumed to originate from a single protoplast, this approach ensures that differences in colony size reflect growth phenotypes rather than variations in the initial amount of starting tissue.

### Gene selection for screening

We utilized publicly available transcriptome data from the Joint Genome Institute (JGI) Gene Atlas Project (http://jgi.doe.gov/doe-jgi-plant-flagship-gene-atlas/), based on *P. patens* genome annotation version 3.3 (Lang *et al*., 2018) and accessed via Phytozome (https://phytozome-next.jgi.doe.gov/) (Perroud *et al*., 2018). A list of known mitotic genes in *P. patens* was compiled based on previous functional studies. This list was not exhaustive but served as a test for our workflow (**Supplemental Table 1**). For each mitotic gene, we manually examined co-expression networks and selected genes with a co-expression score of ≥0.8 (arbitrarily chosen). We further refined this list by excluding genes with previously reported functions (verified through *P. patens* and *A. thaliana* homolog searches) and those with clearly defined non-mitotic roles based on PANTHER annotations (e.g., metabolic enzymes). The final selection of genes is provided in **Supplemental Table 1**.

### *In silico* prediction of protein function and localization

We mapped gene models from the *P. patens* annotation version 3.3 (Lang *et al*., 2018) to the most recent version 6.1 (Bi *et al*., 2024) and found no changes on CDS and thus amino acid level. Annotation of functional domains and possible family associations for our candidate sequences was inferred with InterProScan (version 5.71-102.0; (Jones *et al*., 2014)). We then used DeepLoc (version 2.1; (Ødum *et al*., 2024)), LOCALIZER (version 1.0.4; (Sperschneider *et al*., 2017)), TargetP (version 2.0; (Armenteros *et al*., 2019)) and SignalP (version 6.0; (Teufel *et al*., 2022)) to predict subcellular localization.

### gRNA library construction and molecular cloning

CRISPR gRNAs were designed using the online tool CRISPOR (http://crispor.tefor.net/), based on target gene specificity (off-target score) and predicted frameshift efficiency (Concordet & Haeussler, 2018). The target site was selected within an exon common to all transcript variants per Phytozome data. When multiple homologs were present, we prioritized designing a single gRNA targeting a conserved site. gRNAs were synthesized as single-stranded primers, which also contained overhangs, matching the *Bsa*I-generated overhangs in the destination vector. Forward and reverse primers for each gRNA were mixed together and annealed in the PCR thermocycler. Next, generated individual gRNAs were ligated into pCasGuide/pUC18 vector pre-digested with *Bsa*I, between the U6 promoter and gRNA scaffold sequence (Collonnier *et al*., 2017). The destination vector for gRNA also contained a hygromycin resistance cassette for plant selection. Each plasmid in the gRNA library was verified by sequencing to confirm correct integration of the gRNA. In case when three or more gRNA were required to target all homologs in the respective gene family, we carried out an additional cloning step to assemble all gRNAs into a single vector (Leong *et al*., 2018). In this case, individual gRNAs together with the *U6* promoter and gRNA scaffold region were amplified by PCR and assembled into a single multi-gRNA plasmid using an in-house Gibson assembly mix (Gibson *et al*., 2009).

Plasmids for mNeon-Green endogenous tagging were also assembled using Gibson assembly, with mNeonGreen gene and antibiotic resistance cassette flanked by homologous recombination regions (500–800 bp of the respective genes). A detailed protocol for endogenous gene tagging and knockouts in *P. patens* has been published previously (Yamada *et al*., 2016). RNAi vectors were cloned using the Gateway system (Invitrogen, Carlsbad, CA, USA), with pGG624 as the destination vector (Miki *et al*., 2016). Two independent, non-overlapping RNAi constructs were prepared for each gene. The full list of plasmids and primers used in this study is shown in **Supplemental Table 3**, except for gRNA library primers and plasmids which are shown in **Supplemental Table 1**.

### Microscopy and data analysis

For most samples we used agar-coated glass-bottom dishes for moss culture. A small piece of moss protonema was placed on top of the BCD agar medium and cultured under continuous light at 25 °C for 4-5 days prior to observation (Yamada *et al*., 2016). For RNAi induction, β-estradiol was added to BCD agar medium at a final concentration of 1 □µM (Nakaoka *et al*., 2012). Oryzalin was diluted in 1□mL of distilled water to a final concentration of 20 µM and added directly to the sample during live-cell imaging. For the *cyr1cyr2* frameshift line, due to strong growth defects, we used a microfluidic device to acquire enough data for quantification (Bascom *et al*., 2016). The PDMS device mold was prepared by spin-coating negative photoresist (SU-8 3025) on a silicon wafer, and microchannel designs were created using a maskless lithography system (DL-1000). The PDMS pre-polymer mixture (Sylgard 184) was prepared by mixing the elastomer base and curing agent at a 10:1 ratio, poured onto the mold, degassed for 30 min, and cured in an oven at 65°C for 90 min. The microchannel inlet was punched using a 1.5-mm biopsy punch. The PDMS device and glass-bottom dish were treated with air plasma for 30 seconds, pressed together, and heated at 65°C for 2 h to achieve irreversible bonding. To introduce moss cells in the device, we collected a small piece of protonema grown on cellophane after 5–7 days of culture and blended it in 5 mL BCDAT liquid medium. The suspension was injected into the microdevice. To prevent evaporation, the device was submerged in BCDAT medium, sealed with parafilm, and incubated at 25°C under continuous light for 3–4 days (Kozgunova & Goshima, 2019; Yoshida & Kozgunova, 2023).

Most images were acquired using a Nikon Ti microscope (60 × 1.30-NA lens; Nikon, Tokyo, Japan) equipped with a CSU-X1 spinning-disk confocal unit (Yokogawa, Tokyo, Japan) and an electron-multiplying charge-coupled device camera (ImagEM; Hamamatsu, Hamamatsu, Japan). The microscope was controlled using the NIS-Elements software (Nikon). Imaging after FM4-64 staining and original observation of *cyr1cyr2* frameshift lines (**Supplemental Figure 1B, Supplemental Video 1**) were performed using a ZEISS inverted microscope (25 × 0.8-NA lens or 63 × 1.4-NA lens, Carl Zeiss Microscopy, Germany) equipped with a CSU-X1 spinning-disk confocal unit (Yokogawa, Tokyo, Japan) and CCD camera (Photometrics Prime sCMOS). All imaging was performed at 22–24 °C in the dark. Imaging was performed at least twice for localization and at least three times for phenotypic characterization. To obtain quantitative values, the data from multiple experiments were combined because of insufficient sample numbers in a single experiment. Moss colony images were acquired after 4 weeks of culture using a smartphone camera. Image data were analyzed using ImageJ (National Institutes of Health, Bethesda, MD, USA). Prism was used for the graphs and statistical analyses (GraphPad, San Diego, CA, USA).

### Homolog identification and phylogenetic reconstruction

Based on the Physcomitrella reference sequences identified in this study, we conducted proteome-wide sequence searches with blastp (version 2.16.0+; (Altschul *et al*., 1997)) against *Anthoceros angustus* (Zhang *et al*., 2020b), *Arabidopsis thaliana* (Cheng *et al*., 2017), *Calohypnum plumiforme* (Mao *et al*., 2020), *Ceratodon purpureus* (Carey *et al*., 2021), *Ceratopteris richardii* (Marchant *et al*., 2022), *Chara braunii* (Nishiyama *et al*., 2018), *Chlamydomonas reinhardtii* (Merchant *et al*., 2007), *Chlorella* sp. (Hamada *et al*., 2018), *Cyanophora paradoxa* (Price *et al*., 2019), *Diphasiastrum complanatum* (v3.1, DOE-JGI, http://phytozome-next.jgi.doe.gov), *Dunaliella salina* (Polle *et al*., 2017), *Funaria hygrometrica* (Kirbis *et al*., 2025), *Galdieria sulphuraria* (Rossoni *et al*., 2019), *Klebsormidium nitens* (Hori *et al*., 2014), *Marchantia polymorpha* (Bowman *et al*., 2017), *Mesotaenium endlicherianum* (Cheng *et al*., 2019), *Microglena* sp. (Ye *et al*., 2022), *Nicotiana benthamiana* (Ranawaka *et al*., 2023), *Oryza sativa* (Ouyang *et al*., 2007), *Picea abies* (Nystedt *et al*., 2013), *Populus trichocarpa* (Tuskan *et al*., 2006), *Selaginella moellendorffii* (Banks *et al*., 2011), *Sphagnum fallax* (Healey *et al*., 2023), *Sphagnum magellanicum* (Healey *et al*., 2023), *Spirogloea muscicola* (Cheng *et al*., 2019), *Takakia lepidozioides* (Hu *et al*., 2023), *Vitis vinifera* (Jaillon *et al*., 2007) and *Volvox carteri* (Prochnik *et al*., 2010). To filter out non-homologous sequences, initial blast hits were manually curated, functional annotation was inferred with InterProScan (version 5.71-102.0; (Jones *et al*., 2014)) and several iterations of multiple sequence alignments and maximum-likelihood tree inferences were performed with MAFFT (version 7.525; (Katoh & Standley, 2013)) and FastTree (version 2.1.11; (Price *et al*., 2010)). During the final round, best-fit models were identified with ModelTest-NG (Darriba *et al*., 2020) and maximum likelihood trees were calculated with RAxML-NG (Kozlov *et al*., 2019) using the ‘JTT+I+G4’ (LACH) and ‘JTT+I+G4+F’ (CYR, SpinMi) models and 1000 bootstrap replicates. Final trees were midpoint-rooted and visualized using R (version 4.3.3, R Core Team, 2024) and the ggtree package (version 3.10.1, (Yu *et al*., 2017)).

### Conflict of interest

The authors declare no competing interests.

## Contributions

E.K. and R.R. designed the research project, K.M., N.v.G. and E.K. performed experiments and analyzed the data, E.K. and R.R. acquired funding, all authors wrote and edited the manuscript.

## Data Statement

All data supporting the findings of this study are available within the manuscript and its Supplemental Information or can be obtained from the corresponding author upon request.

## Acknowledgements

We would like to thank Dr. Moé Yamada for comments on the manuscript, and the Life Imaging Center at the University of Freiburg for microscopic use. This work was funded by Japan Society for the Promotion of Science (KAKENHI) 23K14210 and Research Grant for Young Japanese Researchers from the Nakajima Foundation to EK, German Research Foundation DFG under Germany’s Excellence Strategy (CIBSS – EXC-2189 – Project ID 390939984) to RR. We are grateful for the support by the High Performance and Cloud Computing Group at the Zentrum für Datenverarbeitung of the University of Tübingen, the state of Baden-Württemberg through bwHPC and the German Research Foundation DFG through grant INST 37/935-1 FUGG, as well as the de.NBI Cloud within the German Network for Bioinformatics Infrastructure (de.NBI), ELIXIR-DE (Forschungszentrum Jülich and W-de.NBI-001, W-de.NBI-004, W-de.NBI-008, W-de.NBI-010, W-de.NBI-013, W-de.NBI-014, W-de.NBI-016, W-de.NBI-022). Open Access Funding was enabled and organized by Projekt DEAL.

---

**Supplemental Video 1. Cytokinesis failure was observed in *cyr1cyr2 #1* and *#2* CRISPR lines.** Live-cell imaging was performed in *P. patens* apical caulonemal cells in *cyr1cyr2 #1* and *cyr1cyr2 #2* CRISPR frameshift line expressing GFP-tubulin (green) and H2B-RFP (magenta). Images were acquired every 2 min in a single focal plane. Scale, 10 µm.

**Supplemental Video 2. Multiple phragmoplast formation *cyr1cyr2 #2* CRISPR line and CYR1-Citrine overexpression line.** Live-cell imaging was performed in *P. patens* apical caulonemal cells in *cyr1cyr2 #2* CRISPR frameshift line, expressing GFP-tubulin (left panel, green) and H2B-RFP (left panel, magenta), and CYR1-Citrine (right panel, green) overexpression line, which also expressed mCherry-tubulin (magenta, right panel) as a microtubule marker. Secondary phragmoplast formation is indicated with white arrows. Images were acquired every 2 min as a Z-stack (5 µm in 2.5 µm steps), best focal plane is shown. Scale, 10 µm.

**Supplemental Video 3. Localization of CYR1-Citrine and CYR2-Citrine.** Live-cell imaging was performed in *P. patens* apical caulonemal cells expressing mCherry-tubulin (magenta) and one of the following tagged proteins (green): CYR1-Citrine, CYR2-Citrine or Citrine. Images were acquired every 2 min as a Z-stack (5 µm in 2.5 µm steps), best focal plane is shown. Scale, 10 µm.

**Supplemental Video 4. Chromosome missegregation after inducible RNAi knockdown of LACH.** Representative images of mitotic progression and chromosome missegregation caused by depletion of LACH. RNAi was induced by addition of β-estradiol to the growth medium at final concentration of 1 µM, 5–6 days prior to observation. Images were acquired every 2 min as a Z-stack (5 µm in 2.5 µm steps); best focal plane is shown. Scale, 10 µm.

**Supplemental Video 5. Localization of SpinMi-mNeonGreen.** Live-cell imaging was performed in *P. patens* apical caulonemal cells expressing mCherry-tubulin (magenta) and SpinMi-mNeonGreen (green). Images were acquired every 2 min as a Z-stack (5 µm in 2.5 µm steps), best focal plane is shown. Scale, 10 µm.

**Supplemental Table 1.** Genes and gRNA library used for CRISPR genetic screening. **Supplemental Table 2.** Results of *in silico* prediction of subcellular localization **Supplemental Table 3.** *P. patens* transgenic lines, primers, and vectors used in this study

**Supplemental Figure 1.**
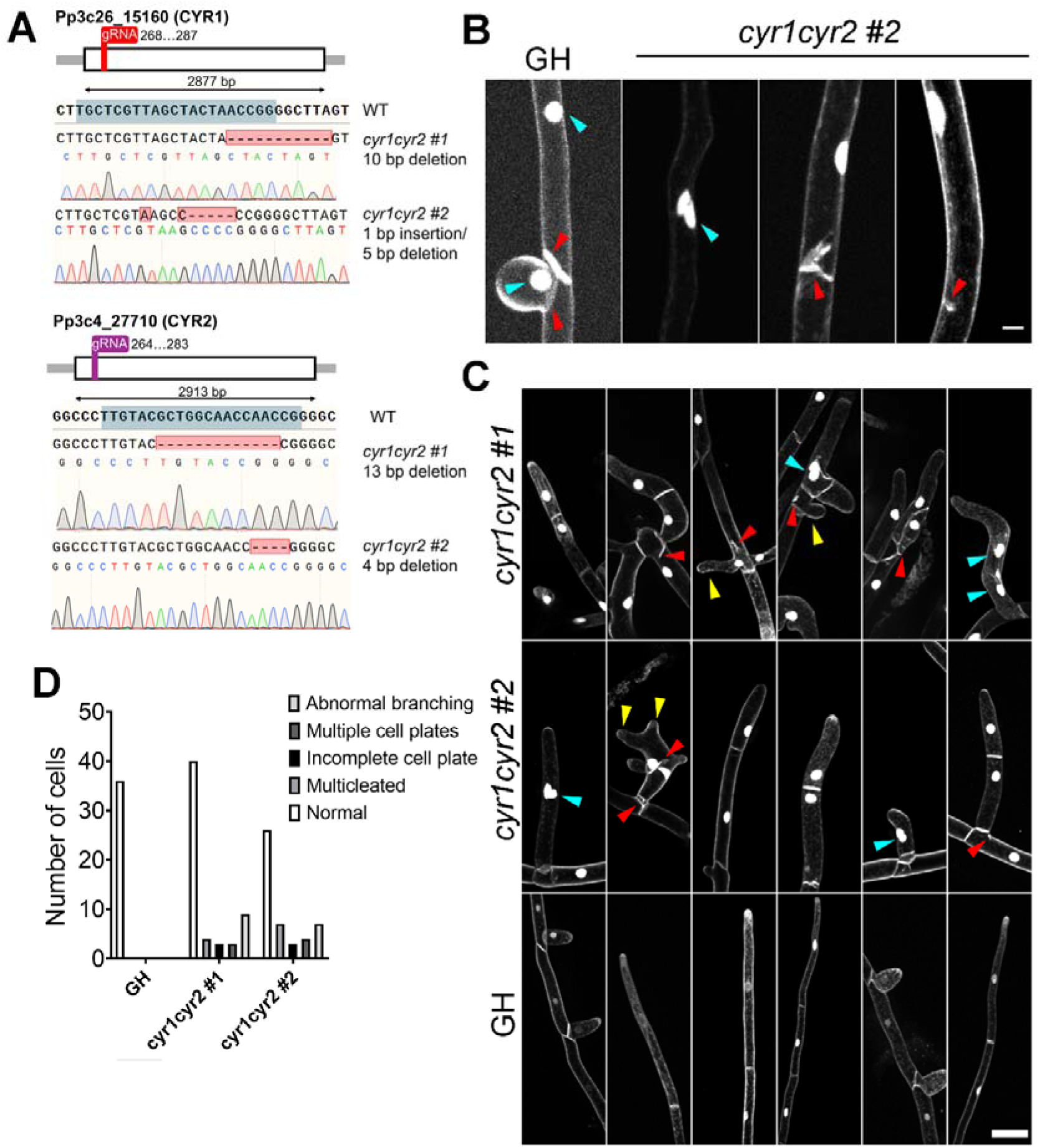
Frameshift verification and initial phenotype observation in *cyr1cyr2#1* and *#2* frameshift lines. (**A**) Gene models for cyr1 and cyr2 genes, and frameshift mutations confirmed by sequencing in *cyr1cyr2 #1* and *cyr1cyr2 #2* moss lines. The gRNA target site is indicated with a blue box. (**B**) Membrane staining with 10 µM of lipophilic dye FM4-64. Nucleus and membranes in same color. Cyan arrowheads point to nuclei, red arrowheads point to cell walls. Note binucleated cell (2^nd^ panel), multiple cell walls (3^rd^ panel), and incomplete cell wall (4^th^ panel) in the *cyr1cyr2 #2* mutant. Scale, 10 µm. **(C)** Membrane staining in the adapted *cyr1cyr2 #1* and *cyr1cyr2 #2* lines after repeated culture. Cyan arrowheads point to multinuclei, red arrowheads point to defective cell walls, yellow arrowheads point to abnormal branching. Scale, 50 µm. **(D)** Number of cells with different phenotypes observed in the adapted *cyr1cyr2 #1* and *cyr1cyr2 #2* lines.

**Supplemental Figure 2.**
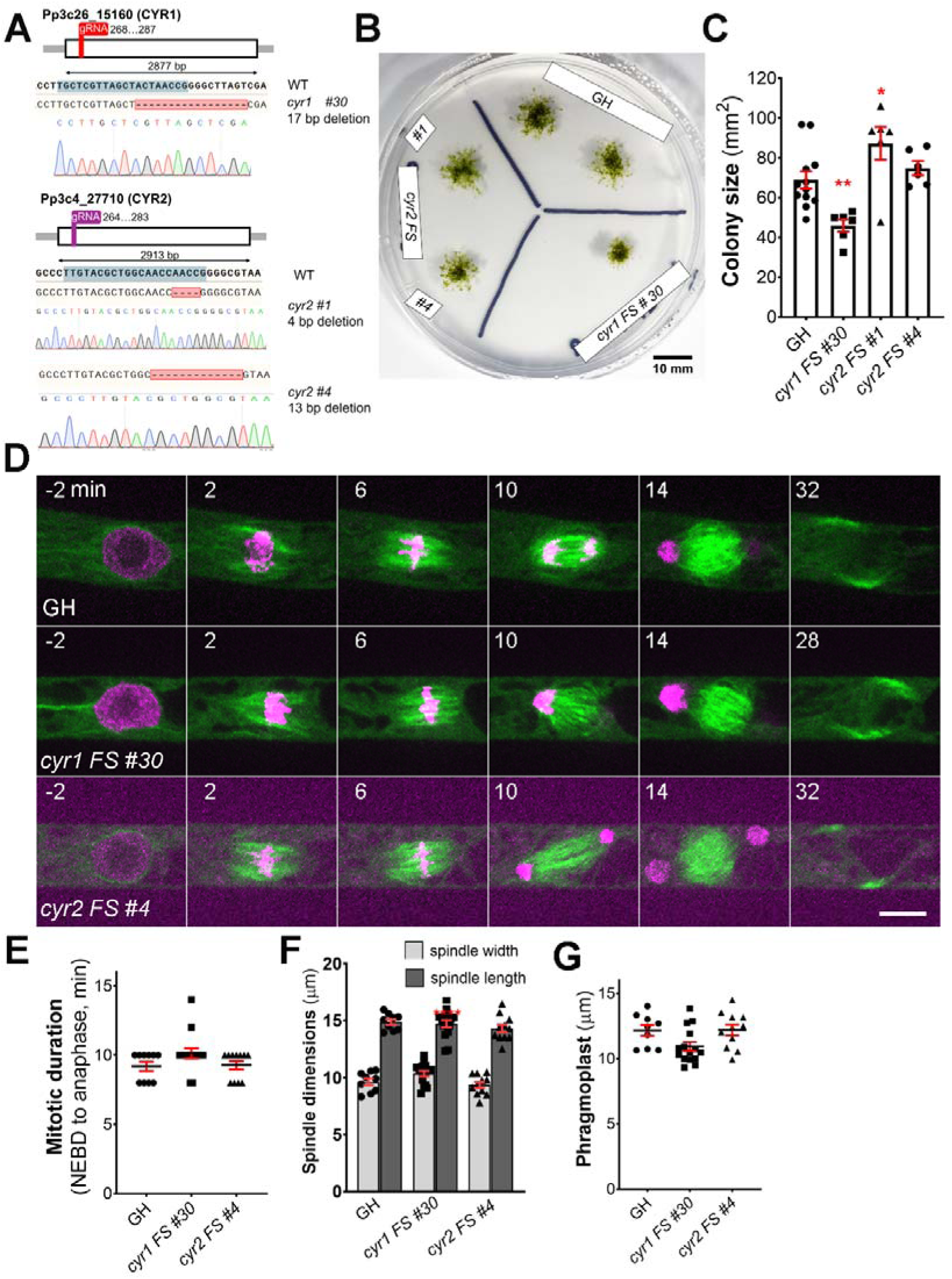
Frameshift verification and phenotype observation in *cyr1* and *cyr2* single frameshift lines. (**A**) Gene models for cyr1 and cyr2 genes, and frameshift mutations confirmed by sequencing in *cyr1#30* and *cyr2 #1* and *#4* lines. The gRNA target site is indicated with a blue box. (B) Representative images of moss colonies after3 weeks of culture. Scale, 10 mm. (C) Colony size measured after 3 weeks of culture. Each data point corresponds to a single colony (mean□±□SEM, **p=0.0089, *p=0.04 by one-way ANOVA with Dunnett’s multiple comparison test against GH) (**D**) Representative images of cell division in GH, *cyr1 FS #30*, and *cyr2 FS #4*. Time zero is set at NEBD, and single focal plane images are shown. Scale, 10 µm. (**E**) Mitotic duration calculated from NEBD to anaphase onset. Each data point corresponds to a single cell (**F**) Spindle dimensions: spindle width (light gray bars) and spindle length (dark gray bars). Each data point corresponds to a single cell (**G**) Phragmoplast length in GH, *cyr1 FS #30*, and *cyr2 FS #4*. Each data point corresponds to a single cell. No statistically significant difference was found in *cyr1 FS #30*, and *cyr2 FS #4* (G-F) against control (GH)

**Supplemental Figure 3.**
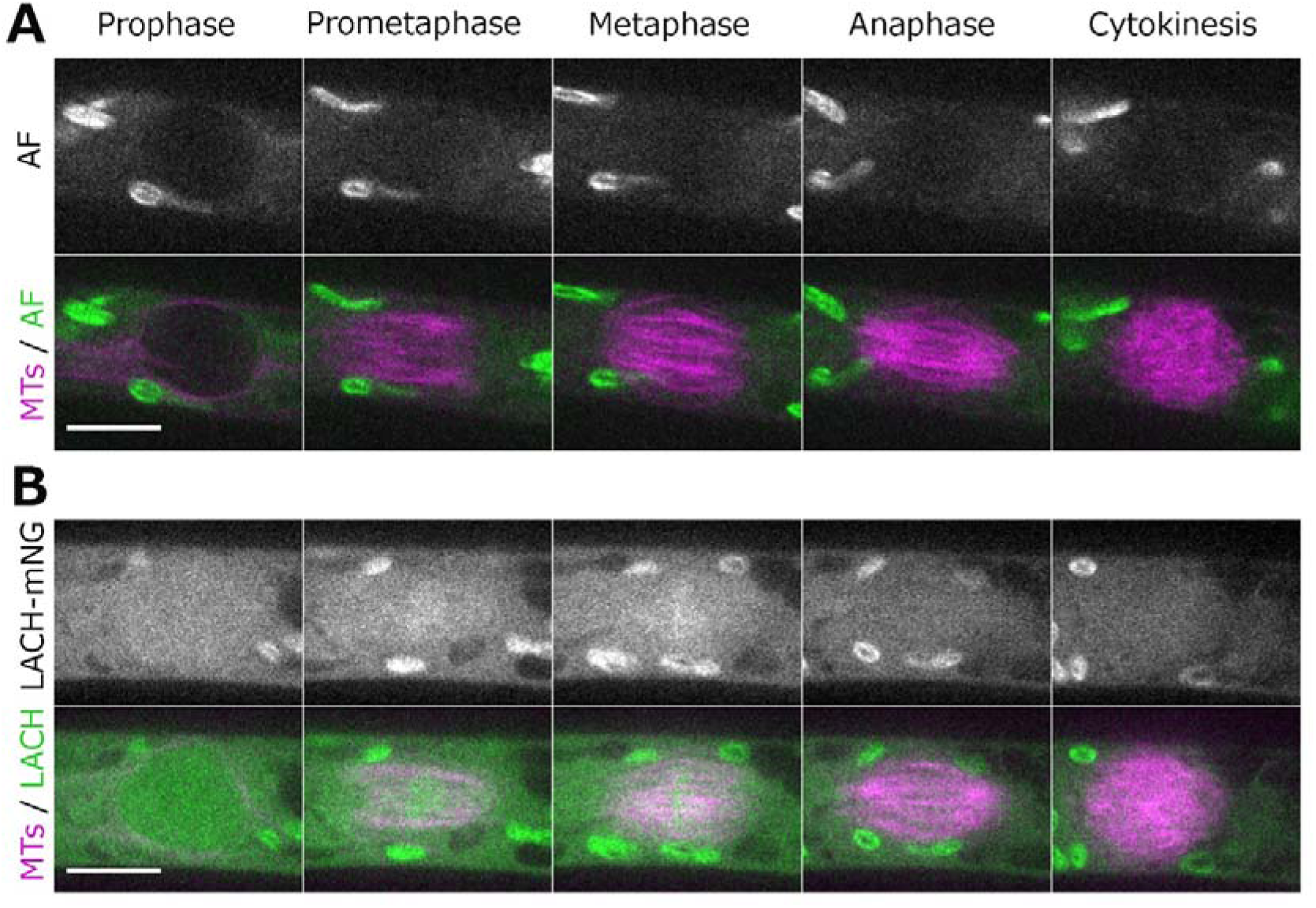
Localization of LACH-mNeonGreen during mitosis. (**A**) Live imaging of mitosis in caulonemal apical cells, expressing mCherry-tubulin as a microtubule marker (magenta). AF panel shows cell autofluorescence background, including chloroplast autofluorescence. Scale, 10 µm. (**B**) Live imaging of *P. patens* protonemal apical cells expressing mCherry-tubulin (magenta) and LACH-mNeonGreen (green). Scale, 10 µm.

**Supplemental Figure 4.**
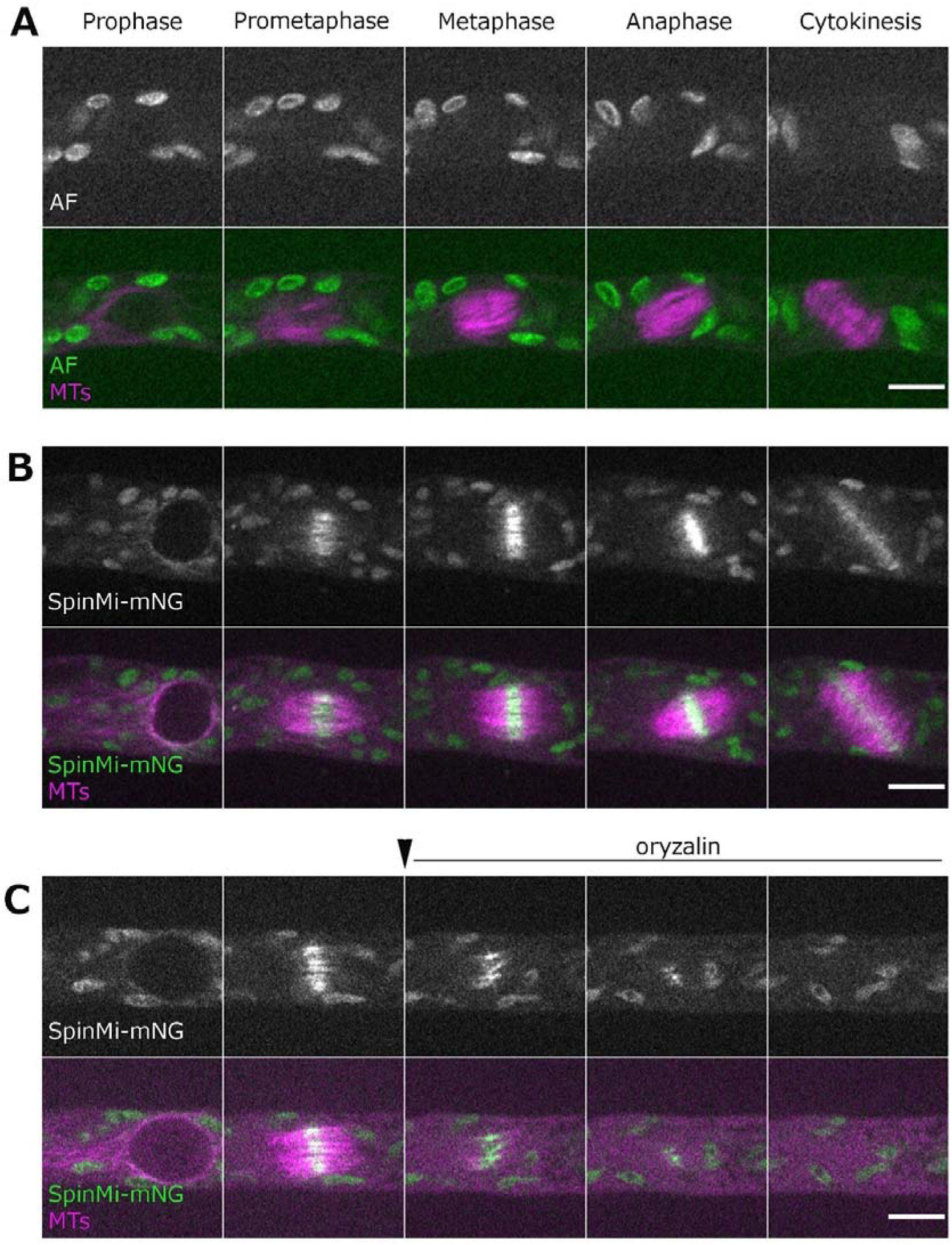
Localization of SpinMi during cell division. Live imaging of *P. patens* caulonemal apical cells expressing (**A**) mCherry-tubulin and (**B**) mCherry-tubulin and SpinMi-mNeonGreen. AF panel shows cell autofluorescence background, including chloroplast autofluorescence. (**C**) Microtubule-depolymerizing drug oryzalin, at final concentration 20 µM, was added during prometaphase to the cell expressing mCherry-tubulin/SpinMi-mNeonGreen. Scale, 10 µm.

**Supplemental Figure 5.**
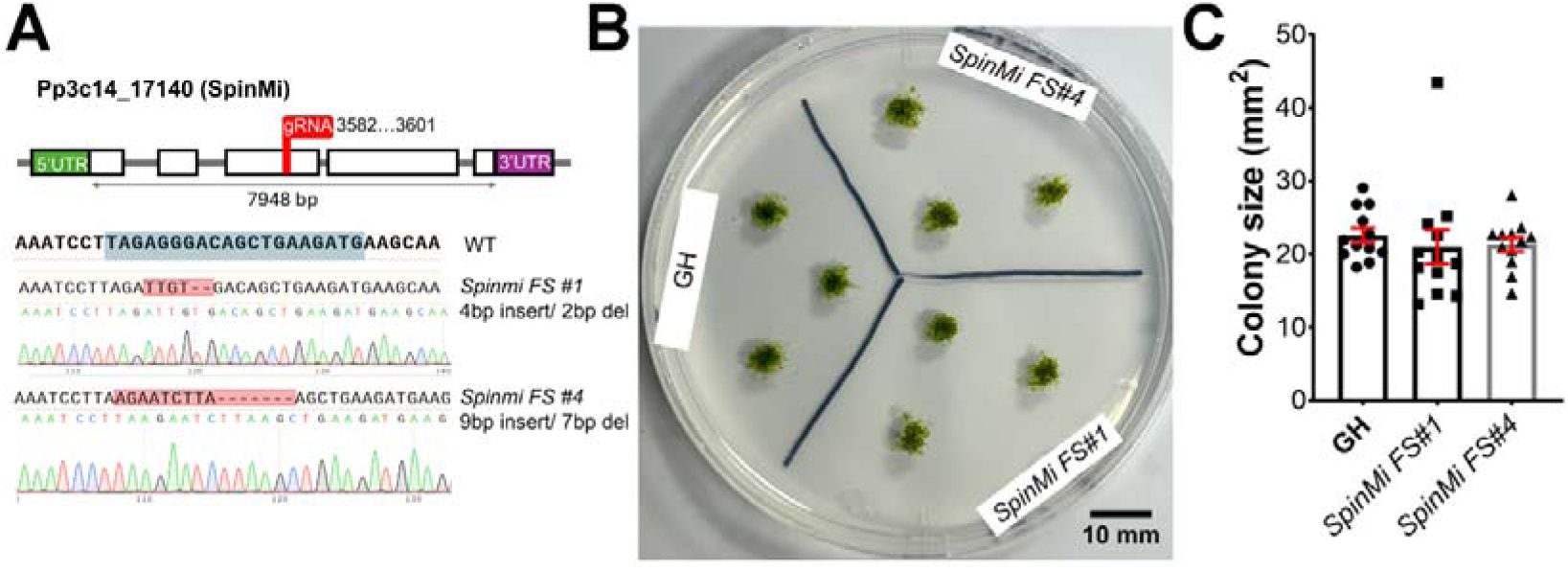
Frameshift mutations in *SpinMi* do not cause serious growth defects. (**A**) Gene model of SpinMi and confirmation of frameshift mutations in the SpinMi gene by sequencing. The gRNA target site is indicated with a blue box. (**B**) Representative images of GH, *SpinMi FS#1* and *SpinMi FS#2* colonies after one month culture. (**C**) Colony size after one month of culture. No statistically significant difference found with one way ANOVA with Dunnett’s multiple comparison test against GH. Each data point corresponds to a single colony (mean±SEM).

## Notes

### Competing Interest Statement

The authors have declared no competing interest.

### Summary of Updates

Added gene models for cyr, spinmi and lach genes, isolated single cyr mutants (Supplemental Figure 2), additional FM4-64 staining (Supplemental Figure 1C, D), and rescue of lach RNAi added (Figure 4)

